# Pcbp1 orchestrates amino acid metabolism burst during the naïve-to-primed pluripotency transition

**DOI:** 10.1101/2025.06.07.658314

**Authors:** E. I. Bakhmet, E. V. Potapenko, O. Y. Shuvalov, A. A. Lobov, E. A. Repkin, N. E. Vorobyeva, A. N. Korablev, A. S. Zinovyeva, A. A. Kuzmin, N. D. Aksenov, A. T. Kopylov, G. Wu, H. R. Schöler, A. N. Tomilin

## Abstract

Embryo implantation is accompanied by the naïve-to-primed pluripotency transition in epiblast cells, making them receptive to external differentiation signals. In addition to this developmental program switch, implantation suggests that an anabolic boost is required for this process, as the embryo-uterine connection begins supplying the requisite nutrients. In this study, we show that the DNA-binding protein Pcbp1 plays a key role in intensifying amino acid metabolism during the priming of pluripotent stem cells. Knockout of the *Pcbp1* gene leads to embryo growth arrest a few days after implantation. By modeling the naïve-to-primed pluripotency transition *in vitro*, we observe reduced proliferation and induction of apoptosis in cells deficient for Pcbp1. Using multi-omics approaches, we uncover a crucial role for Pcbp1 in driving a transcriptional burst of numerous genes involved in the import and the *de novo* synthesis of essential and conditionally essential amino acids. Pcbp1 deficiency is consequently associated with a slowdown in protein biosynthesis, explaining the early lethal phenotype of knockout embryos. Our findings thus uncover the molecular mechanisms underlying anabolic changes during the naïve-to-primed pluripotency transition and highlight the essential role of Pcbp1 in this process, also pointing to its functions in highly proliferative cells.

**Graphical abstract:** 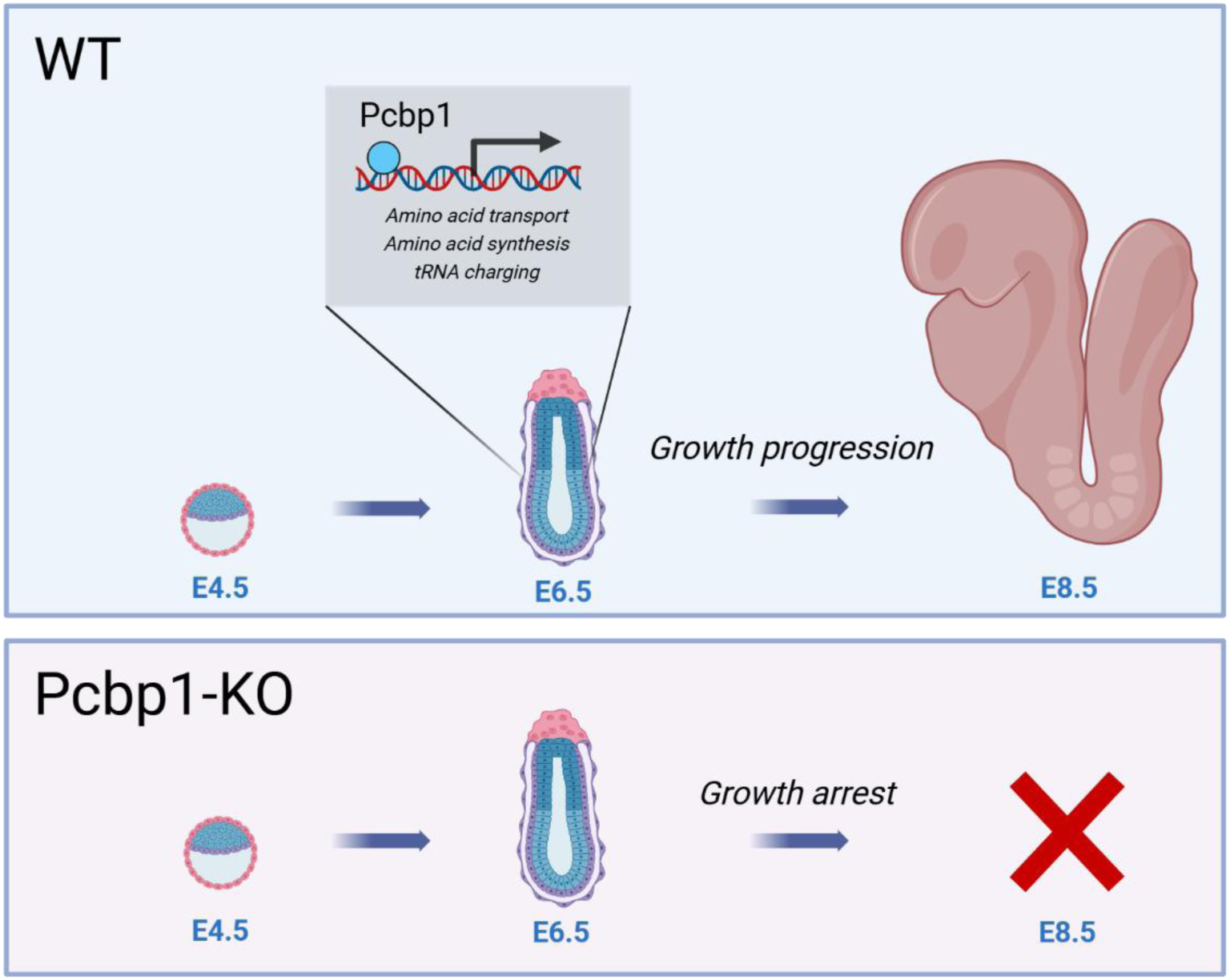

## Introduction

Early mammalian embryogenesis is characterized by a prolonged period without significant body growth. In mice, prenatal development lasts approximately 20 days, with the first five days after fertilization (E5) consisting solely of successive cleavage divisions.

Implantation then establishes the embryo-uterine interaction, providing all essential nutrients for rapid embryo growth. During implantation, pluripotent stem cells of the epiblast undergo the naïve-to-primed pluripotency transition [1–4], a process characterized by changes in the epigenomic [5], gene expression [6, 7], and signaling pathways [8, 9]. At this time epiblast cells become receptive to external differentiation signals [10–12], preparing for gastrulation. While energy metabolism alterations such as enhanced aerobic glycolysis were shown during this phase [13–15], there is still no data describing anabolic changes facilitating rapid embryo growth after implantation.

Pcbp1 is a member of the KH-domain poly(C)-DNA/RNA-binding protein family. This is a ubiquitously expressed protein with a variety of functions including transcriptional regulation [16, 17], mRNA stability [18, 19], alternative splicing [20–23], translation [24, 25], and iron metabolism [26–28]. Several studies indicate its role in erythropoiesis and tumorigenesis, i.e. in highly proliferative cells [23, 29, 30]. Previously, we demonstrated that another KH-domain protein, hnRNP-K, targets open chromatin in mouse naïve embryonic stem cells (ESCs) corresponding to epiblast before implantation but is not required for pluripotency maintenance; rather, it plays a broader role in ESC viability [31]. In turn, ESCs with Pcbp1 knockout are viable [32], though Pcbp1-deficient embryos are not observed few days after implantation [33]. This raises the question about special Pcbp1 functions during the naïve-to-primed pluripotency transition of epiblast cells. Here, we show that growth arrest of Pcbp1 deficient embryos is observed after implantation, while *in vitro* priming of the knockout ESCs results in failed proliferation boost, as well as in cell death by apoptosis. Multi-omics assays unveil crucial role of Pcbp1 in intensification of import and *de novo* synthesis of essential and conditionally essential amino acids during the naïve-to-primed pluripotency transition of epiblast, required for protein biosynthesis and therefore, intensive embryonic growth after implantation.

## Results

### Pcbp1 loss leads to embryonic growth arrest after implantation

To investigate the role of Pcbp1 in early development, we first sought to determine why Pcbp1-knockout (KO) embryos exhibited peri-implantation lethality [33]. To rule out the possibility that maternal Pcbp1 mRNA rescues pre-implantation development, we knocked it down (along with its embryonic counterpart) by injecting Pcbp1 siRNAs into oocytes (Fig. 1A) of the OG2 mice harboring an *Oct4*-*EGFP* transgene [34]. After culturing, we observed that embryos injected with Pcbp1 siRNA developed into blastocysts at a similar rate as those injected with control siRNA (Fig. 1B). RT-PCR analysis confirmed efficient Pcbp1 mRNA depletion (Fig. 1C). Notably, both Oct4 and EGFP expression is properly initiated in Pcbp1-knockdown blastocysts, even though Pcbp1 is known to occupy *Oct4* regulatory elements [35]. In sum, these findings indicate that Pcbp1 is dispensable for both Oct4 expression onset and pre-implantation development.

**Figure 1.**
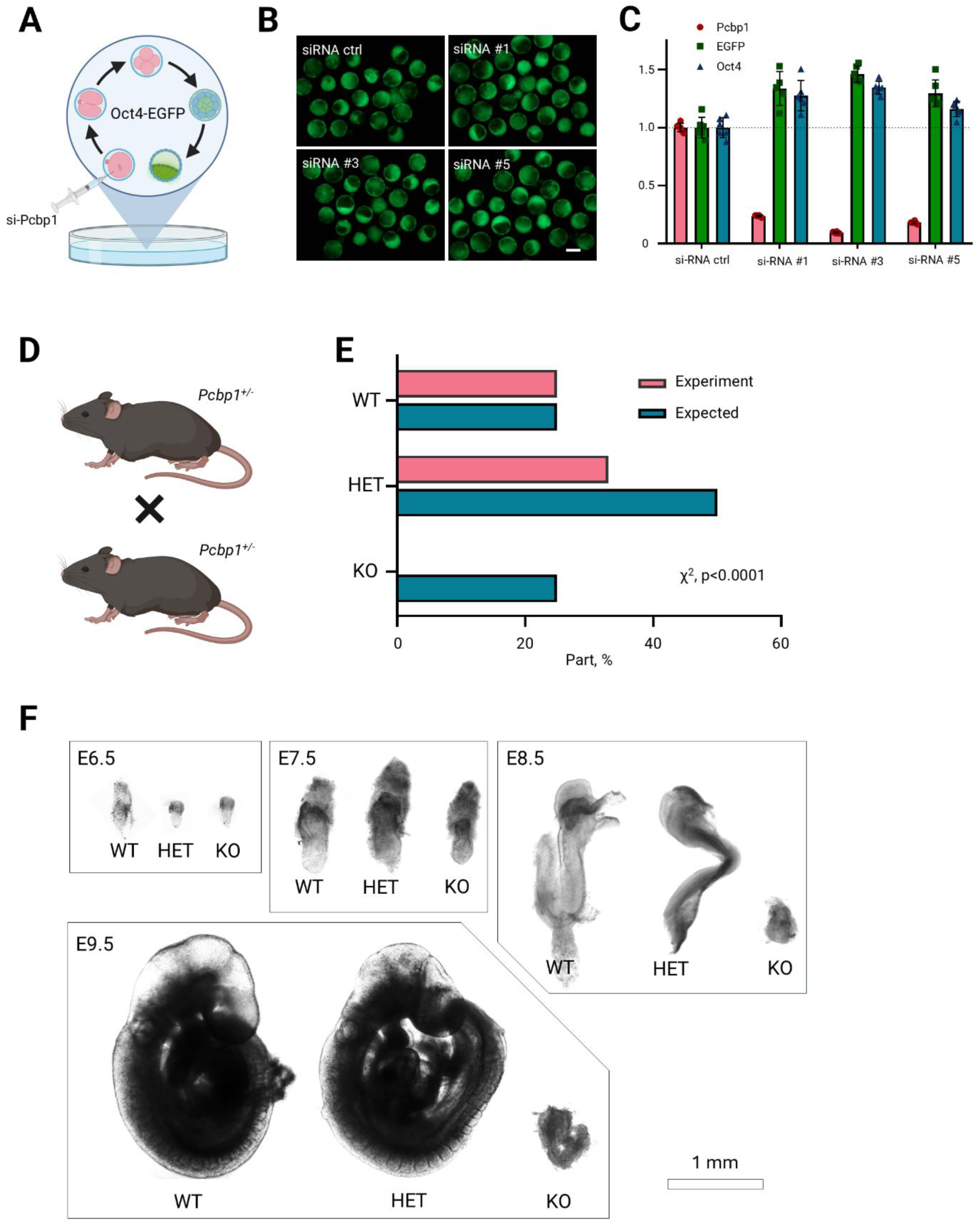
Loss of Pcbp1 function leads to embryo lethality due to impaired post-implantation growth. (A) Schematic representation of the experimental design for Pcbp1 mRNA knockdown in pre-implantation OG2 (Oct4-GFP) embryos. The figure is generated using biorender.com. (B) Fluorescent imaging of OG2 blastocysts derived from zygotes injected with either Pcbp1 mRNA-targeting siRNAs (#1, 3, 5) or control siRNA (ctrl). Scale bar: 100 μm. (C) TaqMan PCR analysis of the blastocysts shown in (B), displaying normalized expression levels of the indicated mRNAs. Data represents 2 biological and 3 technical replicates. (D) Schematic representation of *Pcbp1* heterozygous mice crossbreeding (generated using BioRender.com). (E) Observed and expected percentages of wild-type (WT), heterozygous (HET), and homozygous (KO) *Pcbp1* pups at 2 weeks of age following intercrossing of HET mice. A chi-square statistical test revealed a significant difference between expected and observed ratios (N = 115; p<0.0001). (F) Microphotographs demonstrating a striking difference in embryo size between KO and WT/HET embryos from E6.5 onward.

Next, we used CRISPR/Cas9-mediated gene targeting to generate *Pcbp1*-heterozygous (HET) mice carrying indels at the beginning of the protein-coding sequence (Fig. 1D, Suppl. Fig. 1A, B). Consistent with previous findings [33], genotyping of 2-week–old pups from HET intercrosses revealed no KO offspring and a reduced frequency of HET mice compared to expected Mendelian ratios (Fig. 1E). To further examine the effects of Pcbp1 loss, we performed morphological analysis of post-implantation (E6.5-E9.5) embryos from HET intercrosses (Fig. 1F, Suppl. Fig. 1C). While WT and HET embryos exhibited a progressive size increase, KO embryos demonstrated complete growth arrest from E6.5 onward with the absence of morphologically distinct features.

These findings demonstrate that Pcbp1 loss does not affect pre-implantation development. However, Pcbp1-deficient embryos exhibit an early lethal phenotype, likely due to severe growth defects following implantation.

### Pcbp1 knockout disrupts the naïve-to-primed pluripotency transition in ESCs

We hypothesized that the observed Pcbp1 KO phenotype results from defects in the naïve-to-primed pluripotency transition occurring at and shortly after implantation. To investigate this, we utilized a well-established model recapitulating this transition *in vitro* and relying on the conversion of ESCs into epiblast stem cells (EpiSCs) via the intermediate epiblast-like stem cell (EpiLC) stage (Fig. 2A) [36, 37]. To this end, we used KO Pcbp1 and control (Scr) ESCs, previously generated using CRISPR/Cas9 [32]. No visible differences between Scr and KO cells were observed until Day 2, corresponding to the formative EpiLC stage. However, by Day 4, KO cells exhibited increased cell death and aggregation (Fig. 2B). Timelapse microscopy of Scr cells seeded at low density and cultured under priming conditions showed monolayer formation by Day 4, whereas KO cells demonstrated near-complete proliferation arrest at this stage, followed by partial proliferation recovery (Suppl. Video). Proliferation analysis confirmed that compared to Scr ESCs, KO cells exhibited a reduced growth rate even in the naïve state, (Fig. 2C), which became more pronounced by Day 4—at which stage Scr cells underwent a proliferation burst while KO cells only reached proliferation rates typical of Scr ESCs (Fig. 2C). Surprisingly, there were no differences between Scr and KO cells in G0/G1, S, and G2/M cell cycle phase distribution on Days 0 and 2, suggesting a general cell growth deceleration rather than cell cycle arrest (Suppl. Fig. 2A). However, propidium iodide staining confirmed an increase in KO cell death on Day 4, reaching 20% of the population (Fig. 2D), while annexin V staining revealed that these cells underwent apoptosis (Fig. 2E). Despite these perturbations, KO cells in different pluripotency states showed normal expression of the corresponding markers: ESCs expressed Oct4 and Nanog, EpiLCs expressed Oct4 and Oct6, and EpiSCs were positive for all three markers (Fig. 2F). Western blot analysis of ESC and EpiLC lysates confirmed these results and also showed a significant upregulation of Pcbp1 in Scr cells during the formative pluripotency state (Suppl. Fig. 2B). Consistent with our previous teratoma analysis [32], KO cells retained the ability to give rise to derivatives of ectoderm, endoderm, and mesoderm *in vitro* (Fig. 2G.

**Figure 2.**
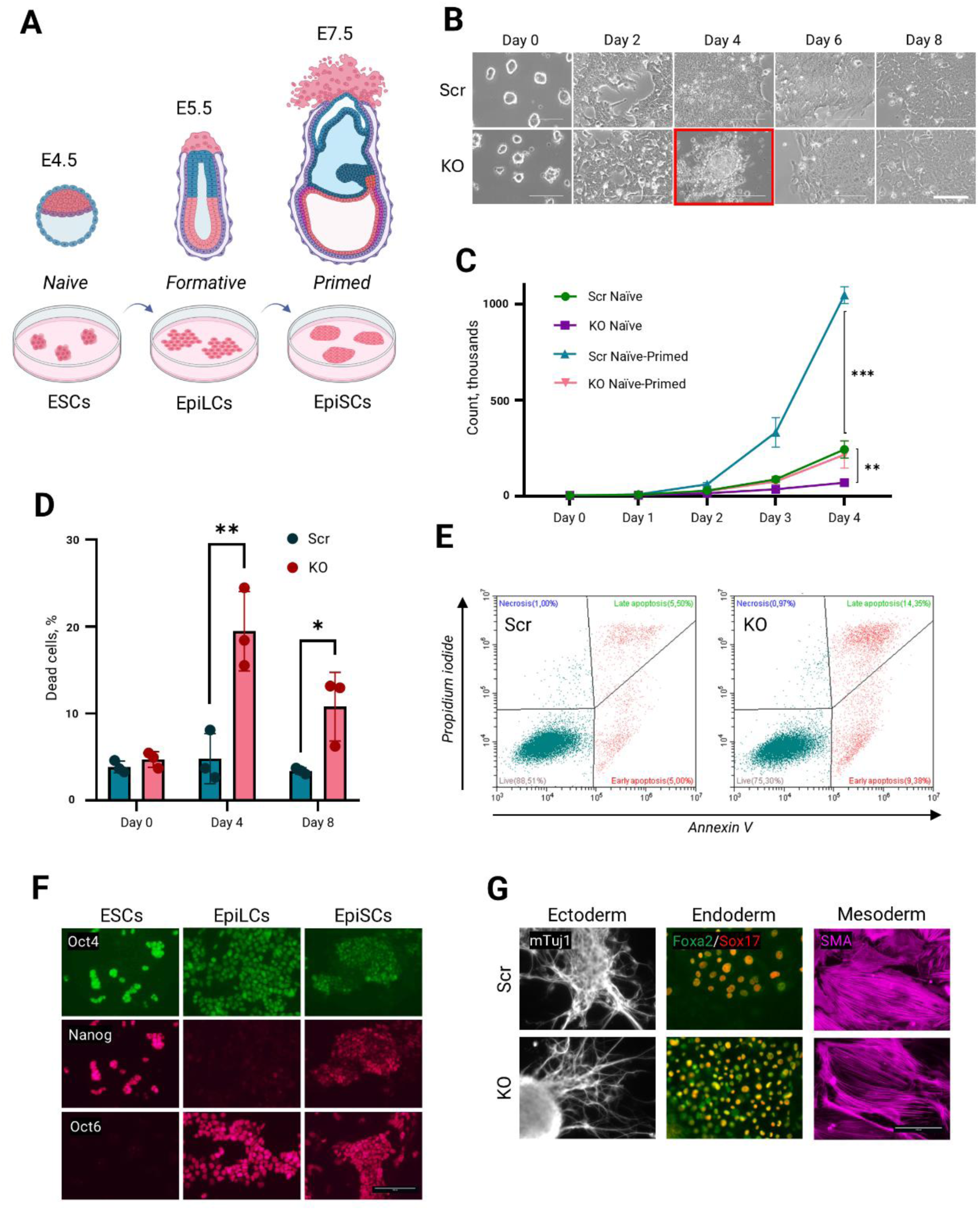
Pcbp1 is essential for the proliferation burst and cell viability during the naïve-to-primed pluripotency transition. (A) Illustration of murine embryos at peri-implantation–stage embryos aligned with corresponding stages of cultured pluripotent stem cells (generated using biorender.com). (B) Microphotographs of Scr and KO cells undergoing the naïve-to-primed pluripotency transition. The red rectangle highlights aggregated and dying KO cells on Day 4, a defining feature of these cells. Scale bar: 100 μm. (C) Proliferation rates of Scr and KO cells in naïve culture conditions and during the naïve-to-primed pluripotency transition. N = 3 biological replicates (individual clones); ** p< 0.01; ****p<0.0001. (D) Propidium iodide (PI) staining to identify dead cells in the naïve state (Day 0) and during the naïve-to-primed pluripotency transition (Days 4 and 8). N = 3 biological replicates (individual cell clones); *p<0.05; **p<0.01. (E) Scr and KO cells on Day 4 of transition stained with PI and antibodies against Annexin V to identify apoptotic cells. (F) Immunofluorescence microscopy of *Pcbp1-*KO ESCs, EpiLCs, and EpiSCs stained with Oct4, Nanog, and Oct6 antibodies. Scale bar: 100 μm. (G) *In vitro–*differentiated Scr and KO cells stained for markers of ectoderm (Tuj1—neurons), endoderm (Foxa2/Sox17—definitive endoderm), and mesoderm (αSMA—smooth muscle cells). Scale bar: 100 μm.

In sum, our findings indicate that Pcbp1 is essential for sustaining viability and proliferation during the transition between the pluripotency substates. The data presented below argues that this is due to failed adaptation to changing metabolic requirements.

### Pcbp1 drives the expression burst of genes involved in amino acid and carbohydrate metabolism during the priming of pluripotent stem cells

To uncover the molecular mechanisms affected by Pcbp1 deficiency during the naïve-to-primed pluripotency transition, we applied a multi-omics approach at early stages before visible morphological defects emerged, specifically, on Day 0 (ESCs) and Day 2 (EpiLCs) (Fig. 3A).

**Figure 3.**
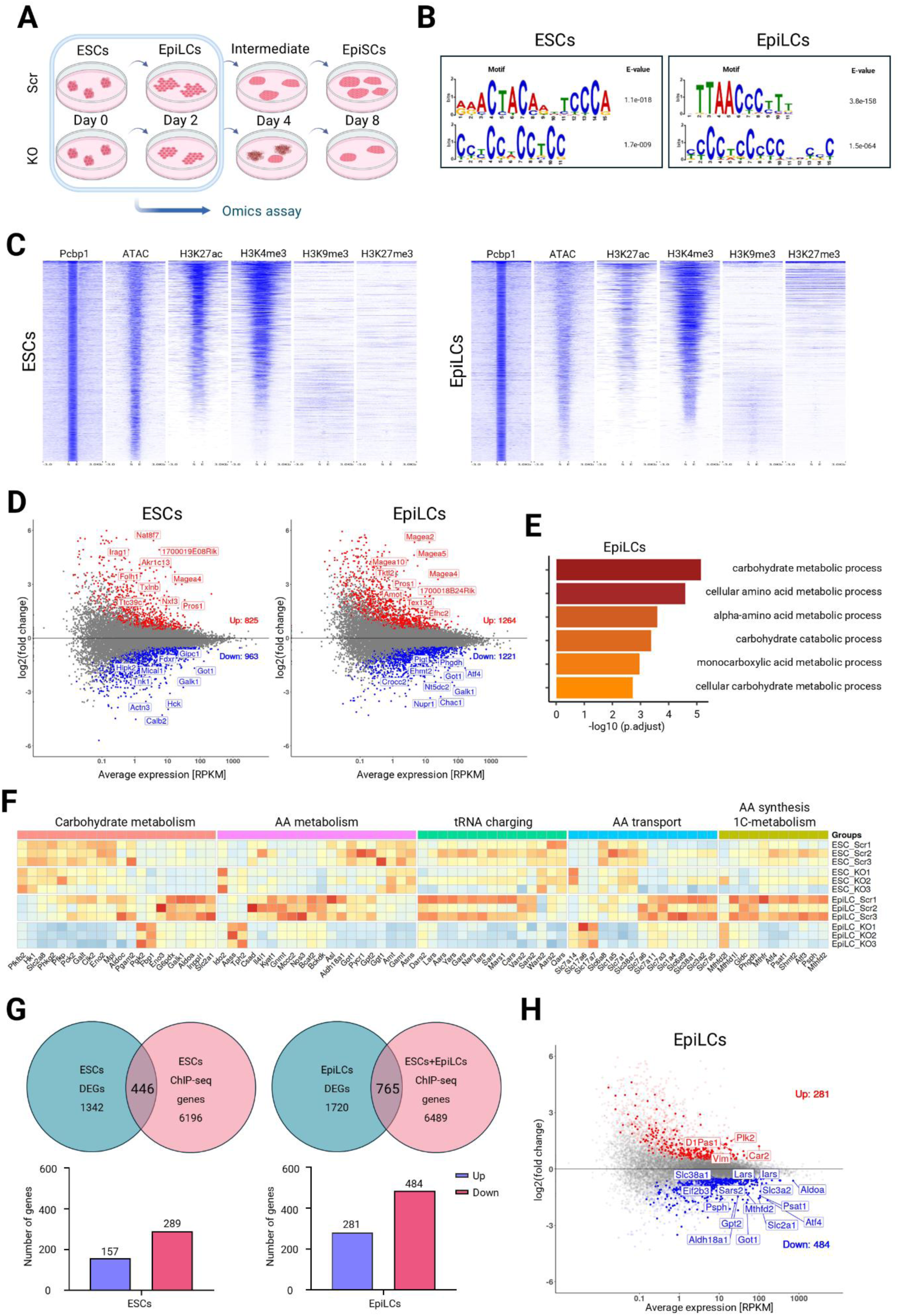
Pcbp1 regulates the expression burst of genes related to carbohydrate and amino acid metabolism in pluripotent stem cells undergoing the naïve-to-primed transition. (A) Schematic representation of timepoints used in omics analyses to identify the molecular mechanisms affected in KO cells. Generated using biorender.com. (B) Pcbp1 *de novo* motifs identified from the ChIP-seq data using the MEME suite at the ESC and EpiLC stages. (C) Heatmaps showing Pcbp1 binding signals in open and transcriptionally active chromatin. ATAC-seq and ChIP-seq data for histone marks in ESCs and EpiLCs were included in the analysis. “S” indicates peak start; “E” indicates peak end. (D) Expression plot of KO differentially expressed genes at ESC and EpiLC stages. The names of the top 20 most significantly differentially expressed genes are highlighted. (E) Gene ontology analysis revealing the top 6 pathways affected in KO cells at the EpiLC stage. (F) Heatmaps showing differentially expressed genes (DEGs) associated with the most affected metabolic processes. (G) Intersection of genes occupied by Pcbp1 in ChIP-seq (+/- 10 kb from TSS) with those differentially expressed in KO cells. ESC DEGs were compared with the ESC ChIP-seq dataset, whereas EpiLC DEGs were compared with both ESC and EpiLC ChIP-seq datasets. (H) Expression plot representing differentially expressed genes with a Pcpb1 binding site nearby (+/- 10 kb, highlighted dots). Semi-transparent dots represent non-occupied differentially expressed genes.

Considering that Pcbp1 is a DNA-binding transcription factor, we first assessed Pcbp1 genome occupancy changes during this transition using ChIP-seq analysis. *De novo* motif search revealed an expected enrichment for poly-C sites in various compositions (Fig. 3B). Previously, we demonstrated that another KH-domain protein, hnRNP-K, binds to open, transcriptionally active chromatin in ESCs [31]. Similarly, Pcbp1 co-localized to open chromatin (ATAC-seq), active enhancers, and transcriptionally active genes (H3K27ac and H3K4me3) in both ESCs and EpiLCs but was absent from heterochromatin regions marked by H3K9me3 and H3K27me3 modifications (Fig. 3C). The vast majority of Pcbp1 binding sites, both in ESCs and EpiLCs, were found within promoters (<1 kb) (Suppl. Fig. 3A) around transcription start sites (TSS) (Suppl. Fig. 3B).

Next, we performed RNA-seq analysis on Scr and KO cells at the ESC and EpiLC stages using three independent Scr and KO clones. Principal component analysis (PCA) demonstrated clear clustering of samples along two primary components—ESCs vs. EpiLCs and Scr vs. KO (Suppl. Fig. 3C). Differential expression analysis revealed 963 downregulated and 825 upregulated genes in ESCs and 1,221 downregulated and 1,264 upregulated genes in EpiLCs (Fig. 3D). Key metabolic genes were significantly downregulated including Atf4, Got1, and Phgdh, which play crucial roles in amino acid metabolism, and Galk1, which is involved in galactose catabolism (Fig. 3D). Consistent with the above findings, no changes in expression were detected for mRNAs associated with naïve pluripotency (Klf4, Nanog, Esrrb, Zfp42, Tbx3) or primed pluripotency (Fgf5, Otx2, Pou3f1, Dnmt3a, Dnmt3b), suggesting that the developmental transition proceeded successfully (Suppl. Fig. 3D). Additionally, we did not detect any notable differences in the expression of genes involved in signaling pathways characteristic of pluripotent stem cells (Suppl. Fig. 3E). Gene ontology (GO) analysis revealed that several metabolic pathways were affected in KO EpiLCs (Fig. 3E). The two most prominently disrupted upon Pcbp1 loss pathways were related to amino acid and carbohydrate metabolism (Fig. 3E). Associated with these pathways genes include those related to glycolysis/gluconeogenesis (Eno2, Eno3, Pfkp, Aldoa, Aldoc, Slc2a1, Galk1, Galt), amino acid metabolism (Got1, Gpt2, Aldh18a1, Nos3, Pycr1), amino acid transport (Slc1a4, Slc3a2, Slc7a3, Slc7a5, Slc7a6), serine/glycine *de novo* synthesis and folate metabolism (Atf4, Phgdh, Psph, Shmt2, Mthfd2, Psat1, Gldc), as well as tRNA charging (Gatb, Iars, Sars, Sars2, Lars, Qars, Vars2) (Fig. 3F). Interestingly, while Scr EpiLCs mostly exhibited significant upregulation of these genes, KO EpiLCs showed no expression change or slight downregulation (Fig. 3F). As expected, the reduced expression of Atf4 in KO EpiLCs (compared to Scr EpiLCs) correlated with reduced expression of known Atf4 target genes, including several regulators of one-carbon (1C) metabolism, amino acid transport, and tRNA charging (Suppl. Fig. 3F).

Next, we investigated genes whose transcription may be directly regulated by Pcbp1. We focused on differentially expressed genes (DEGs) bound by Pcbp1 according to our ChIP-seq data. For this, we cross-referenced ESC DEGs with ESC genes exhibiting Pcbp1 binding within a 10 kb distance from the TSS, while for EpiLCs DEGs, we identified genes occupied by Pcbp1 in both ESCs and EpiLCs (Fig. 3G). We hypothesized that Pcbp1 binding in ESCs could influence gene expression in EpiLCs. The analysis identified 446 potential direct Pcbp1 targets in ESCs and 765 potential direct Pcbp1 targets in EpiLCs. Notably, most of these genes were downregulated in KO cells, suggesting that Pcbp1 functions primarily as a transcriptional activator rather than a repressor (Fig. 3G, lower panel). Among the most notable Pcbp1 transcriptional targets were genes related to amino acid and carbohydrate metabolism, including Atf4, Aldh18a1, Slc38a1, Got1, Sars2, Lars, Mthfd2, Psat1, Aldoa, and Slc2a1 (Fig. 3H, Suppl. Fig. 3G).

Taken together, these results indicate that while KO cells successfully undergo the naïve-to-primed developmental transition, they fail to achieve the necessary amino acid and carbohydrate metabolism intensification during this process. Given the observed genome-wide occupation of Pcpb1, we propose that this metabolic activation is primarily driven by Pcbp1-mediated transcriptional activation of key amino acid and carbohydrate metabolism genes.

### Pcbp1 loss impairs protein biosynthesis during pluripotent stem cell priming

To validate the above findings and assess the physiological state of Pcbp1-deficient cells, we conducted several tests. First, we used a shotgun proteomics approach to analyze protein abundance in Scr and KO cells. As observed in the RNA-seq data, PCA revealed a clear separation between ESCs and EpiLCs, as well as between Scr and KO samples (Fig. 4A).

**Figure 4.**
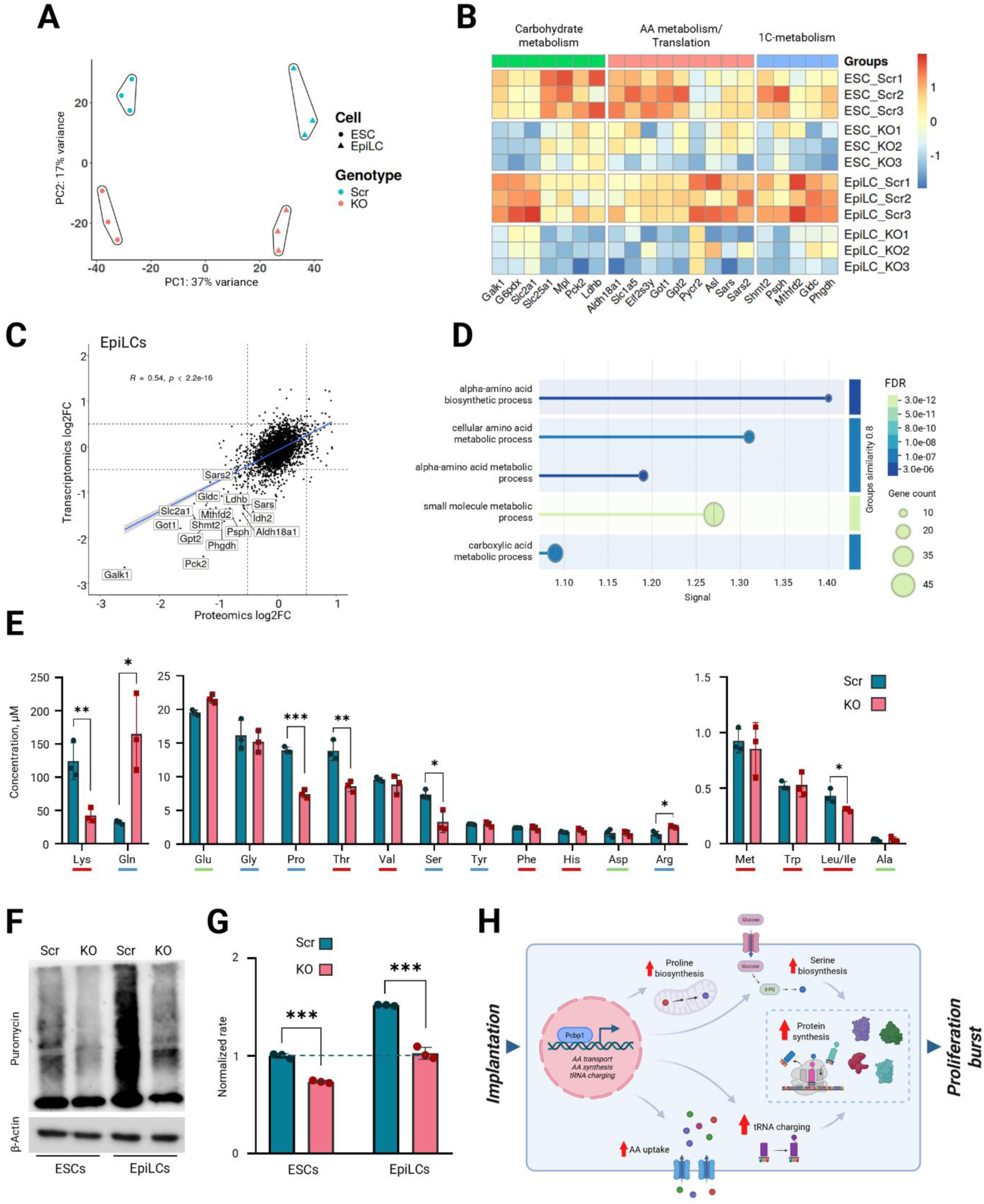
Pcbp1 loss disrupts amino acid metabolism, leading to a decline in protein synthesis during the naive-to-primed pluripotency transition. (A) Principal component analysis showing clustering of samples based on protein abundance comparing Scr vs. KO and ESC vs. EpiLC principal components. (B) Heatmap showing differences in abundance of proteins related to carbohydrate, amino acid, and 1C metabolism (N = 3 biological replicates, i.e. individual cell clones). (C) Intersection of RNA-seq and proteomics analyses, identifying commonly downregulated genes and corresponding proteins in KO EpiLCs. (D) Gene ontology enrichment analysis of differentially abundant proteins in Scr and KO EpiLCs, performed using STRING (string-db.org); proteins with log2FC>0.5 were included in the analysis. (E) Metabolome analysis of free amino acids in Scr and KO EpiLCs. Colored lines represent different amino acids: green – non-essential, blue – conditionally essential, red - essential (N = 3 biological replicates, i.e. individual cell clones); *p<0.05; **p< 0.01; ***p<0.001. (F) SUnSET assay showing differential protein synthesis rates in Scr and KO cells at the ESC and EpiLC stages. (G) Normalized protein synthesis rates in Scr and KO cells at the ESC and EpiLC stages revealed by FACS-based visualization of fluorescently labeled O-Propargyl-puromycin (OP-Puro) incorporation. The dashed line represents Scr ESC protein synthesis rate (N = 3 biological replicates, i.e. individual cell clones). (H) Schematic summary of the results of this study.

Consistent with the RNA-seq data, Scr EpiLCs (compared to Scr ESCs) showed an upregulation of several key metabolic proteins, including Pycr2, Sars2, Shmt2, Mthfd2, Phgdh, related to amino acid metabolism, as well as Galk1 and Slc2a1, related to carbohydrate metabolism (Fig. 4B). In contrast, KO EpiLCs exhibited minimal changes or even downregulation of these proteins. To further correlate RNA-seq and proteomics data, we compared the Log2-Fold Change (FC) of mRNA and protein expression in EpiLCs (Fig. 4C). Most genes identified by the two approaches showed similar Log2-FC values, with no negative correlation (Fig. 4C). As the most notable changes were observed for EpiLCs, we focused further analysis on these cells. GO analysis of differentially expressed proteins highlighted amino acid metabolism as one of the most affected pathways (Fig. 4D). Additionally, a protein-protein interaction network reconstruction via the String database [38] revealed that the central cluster of differentially abundant proteins was enriched for proteins associated with amino acid and carboxylic acid metabolism, with most linked to abnormal survival phenotypes (Suppl. Fig. 4A).

Given the downregulation of genes and proteins involved in carbohydrate and carboxylic acid metabolism, along with the observed proliferation decline in KO cells (Fig. 2C), we hypothesized a possible energy deficiency in these cells. To test this, we measured glycolysis (ECAR) and oxygen consumption rate (OCR, a measure of respiration) using a SeaHorse analyzer in Scr and KO EpiLCs (Suppl. Fig. 4B, C). However, we observed no significant differences in glycolysis, respiration, or calculated ATP production. One possible explanation is that KO cells possess some compensatory mechanisms to align energy metabolism to a reduced protein biosynthesis.

Given the potential disruption in amino acid and carboxylic acid levels in Pcbp1-deficient cells, we performed metabolome analysis. The results showed reduced levels of lysine, proline, serine, leucine/isoleucine, and threonine, along with increased levels of glutamine and arginine (Fig. 4E). Notably, all these amino acids are either essential or conditionally essential. Consistent with the results of energy metabolism (Suppl. Fig. 4B, C), no significant changes were detected in the levels of citrate, succinate, fumarate, malate, lactate, and pyruvate (Suppl. Fig. 4D). These findings suggest that the primary consequence of Pcbp1 loss is dysregulation of amino acid levels, which in turn affects global protein biosynthesis. This conclusion was further supported by two independent methods: Western blot–based SUnSET assay (Fig. 4F) and FACS-based visualization of fluorescently labeled O-Propargyl-puromycin (OP-Puro) incorporation (Fig. 4G). Both approaches demonstrated a reduced protein synthesis levels in both KO ESCs and KO EpiLCs compared to their Scr counterparts. Notably, these levels closely correlated with the proliferation rates of the cells (Fig. 2C), as Scr EpiLCs exhibited an intensification of protein synthesis, whereas KO EpiLCs reached the levels typical of Scr ESCs.

Taken together, our results identify Pcbp1 as a critical regulator of amino acid metabolism intensification – the process which in turn provides protein biosynthesis boost during the naïve-to-primed pluripotency transition. Eventually, this anabolic switch underlies an extensive embryo growth after implantation.

## Discussion

While the naïve-to-primed pluripotency program switch occurring at and shortly after implantation [3, 4, 6, 8, 39, 40], including the shift in energy metabolism preference to aerobic glycolysis [13, 14], is well studied, little was still known about the rearrangement of anabolic metabolism during this development process. In this study, we link, for the first time, the poly(C)-binding protein Pcbp1 to amino acid metabolism in pluripotent stem cells during the naïve-to-primed pluripotency transition, providing insight into why Pcbp1-deficient embryos die shortly after implantation.

We present clear evidence of Pcbp1’s role in positively regulating numerous genes associated with amino acid metabolism during the naïve-to-primed pluripotency transition. A deficiency in several amino acids, along with downregulation of several tRNA synthetases, appears to lead to a slowdown in protein synthesis, followed by decline in proliferation rates. Only essential or conditionally essential amino acid levels are dysregulated in Pcbp1-deficient cells, which aligns with the downregulation of genes encoding key transporters, including Slc1a4, Slc7a1/3/5/6, and Slc3a2. Elevated glutamine and arginine levels, along with decreased leucine/isoleucine and lysine, may be related to the reduced antiporter activity of Slc7a5/6 and Slc3a2 [41]. The low level of proline appears to be due to the downregulation of Slc38a1 [42], as well as the key enzymes responsible for proline synthesis—Aldh18a1 and Pycr [43]. Among DEGs, we also observed Shmt2, Mthfd2, Psat1, Psph, and Phgdh—genes related to one-carbon (1C) metabolism and serine biosynthesis. These metabolic processes, which are crucial for both proliferation and amino acid metabolism, are typically upregulated in proliferating cells [44–46]. 1C metabolism relies on glucose as a substrate, which may explain the downregulation of glycolytic genes despite no detected changes in energy metabolism. Additionally, we demonstrate that Atf4, which transcriptionally regulates all the aforementioned processes—from amino acid uptake and biosynthesis to tRNA charging [47–50]—is positively regulated by Pcbp1.

Interestingly, our findings suggest that Pcbp1 functions upstream of Atf4 in the regulatory hierarchy controlling amino acid metabolism, as both Atf4 and its targets are downregulated in KO EpiLCs. Moreover, unlike Pcbp1 KO embryos, Atf4-deficient mice are viable, though they are born with several abnormalities [51]. The lethal phenotypes observed in embryos with knockouts of amino acid transporters [52, 53], tRNA synthetases [54], and 1C metabolism genes [55] further suggest that Atf4 is not essential for mammalian embryogenesis.

Notably, Atf4 [51, 56, 57], along with Pcbp1 and Pcbp2 [29, 33], has been shown to be important for definitive hematopoiesis. Conditional EpoR-driven Cre-mediated knockout of both Pcbp1 and Pcbp2 results in the absence of liver erythropoiesis at E12.5 and embryonic lethality at E13.5 [29]. Interestingly, the depletion of either Pcbp1 or Pcbp2 alone does not lead to lethality in this system, highlighting a unique and essential role for Pcbp1 in the naïve-to-primed pluripotency transition. We propose that both of these processes, which are characterized by high proliferation rates [58], require intensified amino acid metabolism, a function dependent on Pcbp1.

Some findings also suggest a positive role for Pcbp1 in tumorigenesis. Extensive research has been conducted on the tumor-related functions of Atf4 [59–61], Slc1a5 [62], Slc3a2 [63], Slc7a5 [64], as well as on serine [65] and proline biosynthesis [66, 67] in cancer. However, reports on Pcbp1’s role in carcinogenesis are conflicting. While some studies indicate that Pcbp1 functions as a tumor suppressor [22, 68–72], several articles describe its pro-tumorigenic functions via mRNA stabilization [73], TGFβ-signal transduction [23], or ferroptosis inhibition [30, 74]. However, no studies to date have linked Pcbp1 functions in amino acid metabolism to cancer. Interestingly, Atf4 has also been shown to prevent ferroptosis in hepatocellular carcinoma cells [75], yet paradoxically it can also inhibit proliferation and induce apoptosis in tumors [76, 77]. Taken together, these results suggest that Pcbp1’s role in tumorigenesis may be context dependent, and our results highlight the need for further investigation into its role in amino acid metabolism within this process.

Several open questions remain for future research. Surprisingly, as evidenced by SeaHorse analysis, Pcpb1 KO cells were able to adapt their energy metabolism to decreased protein biosynthesis, suggesting the existence of a regulatory loop between these two major metabolic processes. We observed a severe downregulation of both RNA and protein levels of Galk1, Got1, and Gpt2 in Pcbp1 KO cells. While little is known about Galk1’s role in pluripotent stem cells, this enzyme participates in galactose catabolism and may contribute to the production of glucose-6P for serine biosynthesis. Gpt2 catalyzes the conversion of pyruvate and glutamate into alanine and α-KG, while Got1 bidirectionally converts asparagine and α-KG into glutamate and oxaloacetate. Interestingly, despite the loss of Pcbp1 function, we observed no significant changes in pyruvate and TCA metabolites or different levels of glutamate, alanine, and asparagine. This suggests the presence of compensatory mechanisms that balance amino acid levels, possibly through inhibition of glutamine conversion to glutamate to prevent excess glutamate accumulation. Finally, the precise mechanism by which Pcbp1 regulates amino acid metabolism remains to be elucidated. Our data suggests that Pcbp1 plays a role in transcriptional regulation of the genes mentioned above. Recent studies indicate that Pcbp1stabilizes secondary DNA structures known as i-motifs, thereby preventing the formation of G-quadruplexes on the opposite strand [16, 17]. However, other functions of Pcbp1—such as mRNA stabilization, splicing, and translation—should also be examined in the context of pluripotent stem cells.

## Acknowledgements

The study was supported by the Russian Science Foundation (RSF) grant № 23-75-10096, https://rscf.ru/en/project/23-75-10096/. The study was carried out using the equipment of the Employee Facility “Center for Cell Technologies of the Institute of Cytology of the Russian Academy of Sciences” and shared research facility “Vertebrate cell culture collection”.

Schematic images were created using BioRender.com. The UHPLC-MS/MS analysis was conducted using equipment from the Resource Centre for Molecular and Cell Technologies at St. Petersburg State University. We thank Areti Malapetsas for editing the manuscript.

## Contributions

E.I.B. conceived the study, conducted the majority of the experimental procedures, including knockout embryos analysis and cell culture work, wrote the first draft; E.V.P. performed the bioinformatics analysis; O.Y.S. conducted experiments on energy metabolism using a SeaHorse analyzer, measured protein biosynthesis rates, and participated in omics data interpretation; A.A.L. and E.A.R. performed shotgun proteomics and proteomics data analysis, with A.A.L. also contributing to omics data interpretation; N.E.V. prepared DNA libraries for ChIP-seq analysis; A.N.K. established *Pcbp1^+/-^* heterozygous mice; A.S.Z. contributed to cell culture work; A.A.K. assisted with molecular cloning; N.D.A. performed FACS analysis; A.T.K. carried out metabolome analysis; G.W. conducted experiments with pre-implantation embryos; H.R.S. provided guidance throughout the project and contributed to the wring; and A.N.T. conceived the study, managed the project, contributed to materials, tools, and reagents, and edited and approved the final version of the manuscript.

## Methods

### Pcbp1 knockdown in preimplantation embryos

Fertilized oocytes were collected in M2 medium 18 hours post-hCG from the oviducts of primed OG2 female mice after mating with OG2 male mice. The oocytes were cultured in KSOM medium at 37°C in a 5% CO2 atmosphere until microinjection. Three siRNA duplex oligonucleotides targeting Pcbp1 mRNA were tested: siRNA-#1 (5’-GTGTGACTGAAAGCGGACTCA-3’), siRNA-#3 (5’-GAACCAGGTGGCAAGACAA-3’), and siRNA-#5 (5’-GCTGATGCACGGAAAGGAA-3’); siRNA-ctrl (5’-GCACCCGATAAGCGGTCAA-3’) was used as a negative control. The lyophilized siRNA duplexes (Dharmacon, Lafayette, CO) were resuspended in siRNA buffer (Dharmacon, cat. no. B-002000-UB-100) according to the manufacturer’s instructions and stored in single-use aliquots at −20°C.

Microinjection of siRNAs was performed using a FemtoJet microinjector (Eppendorf) and a micromanipulator (Narishige) in M2 medium drops covered with mineral oil.

Microinjection pipettes were pulled using a Sutter P-97 pipette puller. Five microliters of siRNA (20 μM) were loaded into the pipette and approximately 2 pL of siRNA solution were injected into the cytoplasm of each oocyte. Following microinjection, oocytes were washed and cultured in KSOM medium at 37°C in a 5% CO2 atmosphere, with cleavage evaluated twice daily. After 2 days in culture, embryos that had developed to the blastocyst stage (E3.5) were analyzed for mRNA by TaqMan-qPCR. Animal care was conducted in accordance with the institutional guidelines of the Max Planck Institute.

### Generation of *Pcbp1*-heterozygous mice

Guide RNAs (gRNAs) were designed using the Benchling online service (https://benchling.com/) and synthesized as described previously [78]. For intracytoplasmic microinjection, *in vitro–*fertilized oocytes were obtained from superovulated C57BL/6 females (4-6–week old, ICG SB RAS, Novosibirsk) and spermatozoids from C57BL/6 males (3-month old, ICG SB RAS, Novosibirsk) following established protocol [79]. Fertilized oocytes were microinjected with one of two solutions (1) 120 ng/μL ssODN-1, 25 ng/μL sgRNA-1, 25 ng/μL sgRNA-2, and an equimolar concentration of Alt-R HiFi Cas9 Nickase V3 (IDT, Coralville, IA, USA) or (2) 120 ng/μL ssODN-2, 25 ng/μL sgRNA-4, and an equimolar concentration of Alt-R HiFi Cas9 Nuclease V3 (IDT, Coralville, IA, USA). After microinjection, zygotes were cultured overnight at 37°C in a 5% CO2 atmosphere, and resulting 2-cell–stage embryos were transferred into the oviducts of pseudopregnant CD-1 females (8-10 weeks old, ICG SB RAS, Novosibirsk). Offspring were genotyped for target modifications using Pcbp1-gen primers (Table 3), and PCR products were sequenced by the Sanger method. Founders #6 (from experiment with Cas9-nuclease) and #19 (from experiment with Cas9-nickase) carrying loss-of-function *Pcbp1* alleles were selected and maintained through breeding with C57BL/6 mice. Heterozygous Pcbp1 (HET) offspring were intercrossed to generate homozygous (KO), HET, and WT embryos for developmental analysis. Successful mating was confirmed by the presence of a copulation plug, and embryos were staged as embryonic day (E) 0.5 from noon on the day of the plug detection. Post-implantation embryos were dissected from the uterus, placed in PBS, photographed using the EVOS fl Auto visualization system (ThermoFisher), and genotyped using Pcbp1-gen primers (Table 3). All animal procedures and technical manipulations were performed in compliance with the European Communities Council Directive of 24 November 1986 (86/609/EEC) and were approved by the Bioethical Committee at the Institute of Cytology and Genetics (Permission N45 from 16 November 2018).

### Cell culture work

Murine embryonic stem cell (ESC) mutants used in this study were derived from Tg2a E14 ESCs (Bay Genomics). Control cell lines (Scr1, Scr2, Scr3), and *Pcbp1-*knockout lines (KO1, KO3, and KO22) were previously reported [32]. Unless otherwise specified, all reagents for mouse ESC culturing were purchased from Gibco (ThermoFisher Scientific). ESCs were routinely cultured on adhesive plastic dishes (Eppendorf or TPP) precoated with 0.1% gelatin (Merck) under standard conditions (5% CO2, 37°C). The mES medium consisted of Knockout-DMEM supplemented with 15% fetal bovine serum (HyClone), 1x penicillin/streptomycin, 2 mM L-Glutamine, 1x non-essential amino acids, 50 μM β-mercaptoethanol, and 500 U/mL bacterially expressed human LIF produced in-house. Naïve ESC culturing and EpiLC induction were performed as described previously [3]. ESCs were first cultured for 6-8 days on the poly-L-ornithine–coated plastic in 2i-LIF-N2B27 medium. This medium was a 1:1 mixture of N2 (DMEM/F12 + GlutaMAX with addition of N2 supplement, 1x Pen-Strep, 0.005% BSA, and 50 µM β-mercaptoethanol) and B27 (Neurobasal medium with addition of B27 supplement, 2 mM L-Gln, 1x Pen-Strep and 50 µM β-mercaptoethanol), supplemented with 500 U/mL hLIF, 3 µM CHIR99021 (Axon) medium, and 1 µM PD0325901 (Axon). ESCs were passaged using 0.05% Trypsin-EDTA and were used for EpiLC/EpiSC derivation after no more than 8 days in 2i-LIF-N2B27 culture, as prolonged MEK inhibition is known to induce genomic and epigenetic changes [80, 81]. For derivation of EpiLCs, 28,000 naïve ESCs per cm^2^ were seeded on fibronectin (Merck)-coated (15 µg/ml) plastic in EpiLC medium: N2B27 supplemented with 12 ng/mL bFGF (Peprotech), 20 ng/mL Activin A (Peprotech), and 1% knockout serum replacement. For further maturation of EpiSCs [36, 37], XAV939 was added from day 2 (EpiLC stage) to prevent spontaneous differentiation [82, 83]. Cells were passaged using collagenase IV, and 10 µM XAV939 was added to the EpiLC medium. Due to differences in proliferation rates of Scr and KO cells, KO cells were seeded at 2.3-fold the density of Scr cells on Day 0 to ensure equal cell numbers for energy profiling and proteomic, metabolomic, and protein biosynthesis assays.

### Cell death analysis

Propidium iodide (PI) was added directly to the medium at a final concentration of 50 µg/mL for 2 minutes. The medium containing PI and floating cells was collected and combined with cells harvested from the well using Trypsin/EDTA. Cells were centrifuged, washed, and resuspended in PBS before being analyzed by FACS. Alexa Fluor 647–labeled annexin V antibodies were used according to the manufacturer’s recommendations.

### Immunocytochemical analysis

Cells were fixed in 4% paraformaldehyde (Merck) for 10 minutes at room temperature (RT), permeabilized with 0.1% Triton X-100 for 15 minutes, and blocked in 3% BSA for 1 hour at RT. Primary antibody staining was performed overnight at 4°C using the appropriate antibodies (Table 1) diluted in PBS containing 0.1% Tween-20 (PBST). Cells were then washed 5-6 times with PBST, stained with secondary fluorescent antibodies (Jackson ImmunoResearch) for 2 hours at RT, and washed 2 or 3 additional times with PBST. Nuclei were stained with DAPI in PBST for 5 minutes at RT, and cells were stored in PBS containing NaN3 until imaging. Fluorescent microscopy was performed using an EVOS FL AUTO (Life Technologies) microscope equipped with DAPI, GFP, RFP, and CY5 filter cubes.

**Table 1.** Antibodies used in this study.

| Target | Cat. No. | Manufacturer | Application |
| --- | --- | --- | --- |
| Pcbp1 | ab74793 | Abcam | ChIP/WB |
| Oct4 | sc-5279 (C-10) | Santa Cruz | ICC/WB |
| Nanog | A300-397A | Bethyl | ICC |
| Oct6 | ab272925 | Abcam | ICC |
| Foxa2 (HNF-3β) | sc-374375 | Santa Cruz | ICC |
| Sox17 | AF1924 | RD systems | ICC |
| Klf4 | ab129473 | Abcam | WB |
| Tuj1 | MMS-435P | Covance | ICC |
| α-SMA | A2547 | Sigma | ICC |
| Gapdh | 2118 | Cell Signaling | WB |
| β-Actin | 8457 | Cell Signaling | WB |
| Annexin V | A23204 | Invitrogen | Apoptosis |
| Puromycin | MABE 343 | Merck | Protein biosynthesis |

Secondary fluorescent and HRP-conjugated antibodies were purchased from Jackson ImmunoResearch.

**Table 2.**
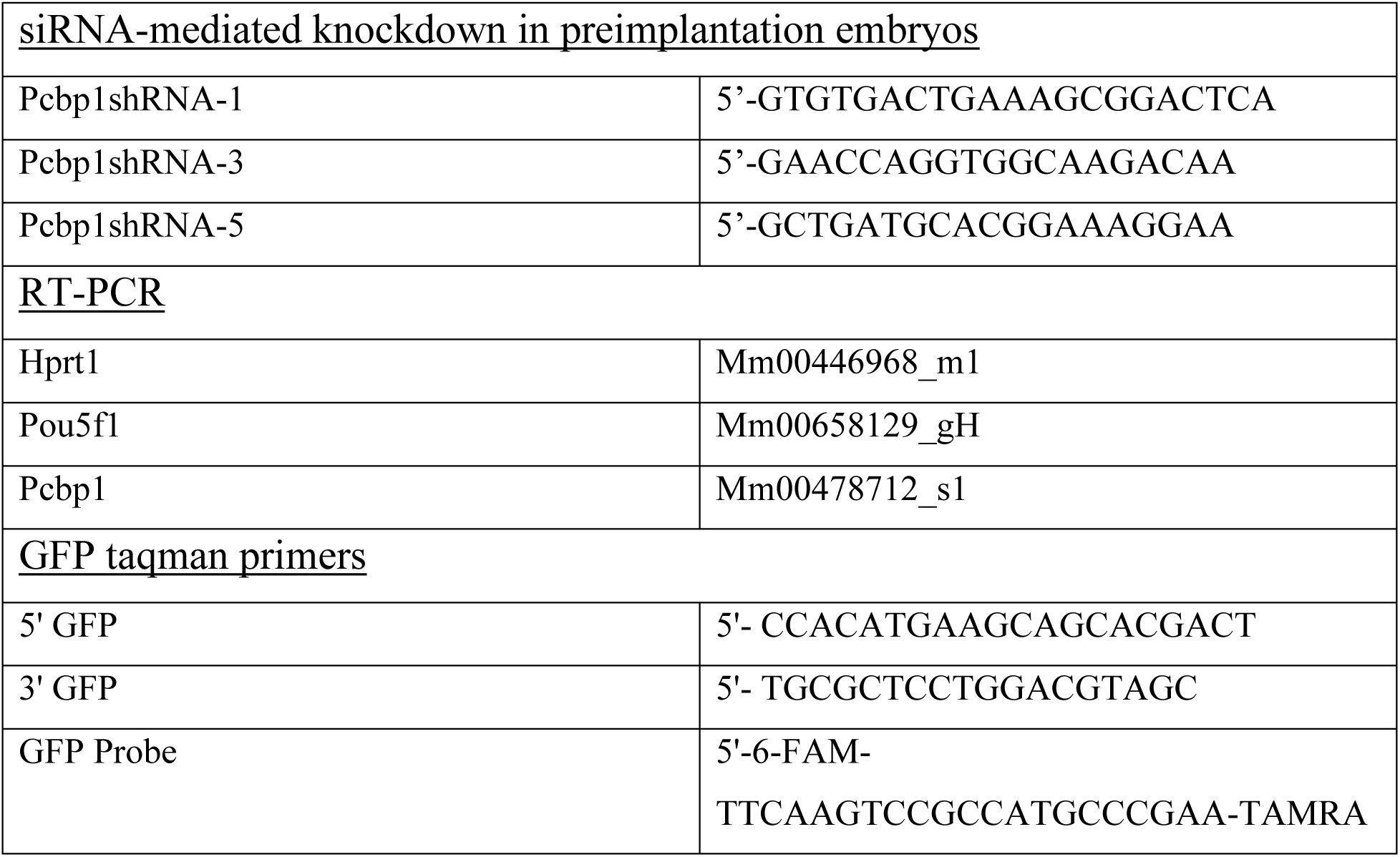
Oligonucleotides used for pre-implantation embryo analyses.

**Table 3.**
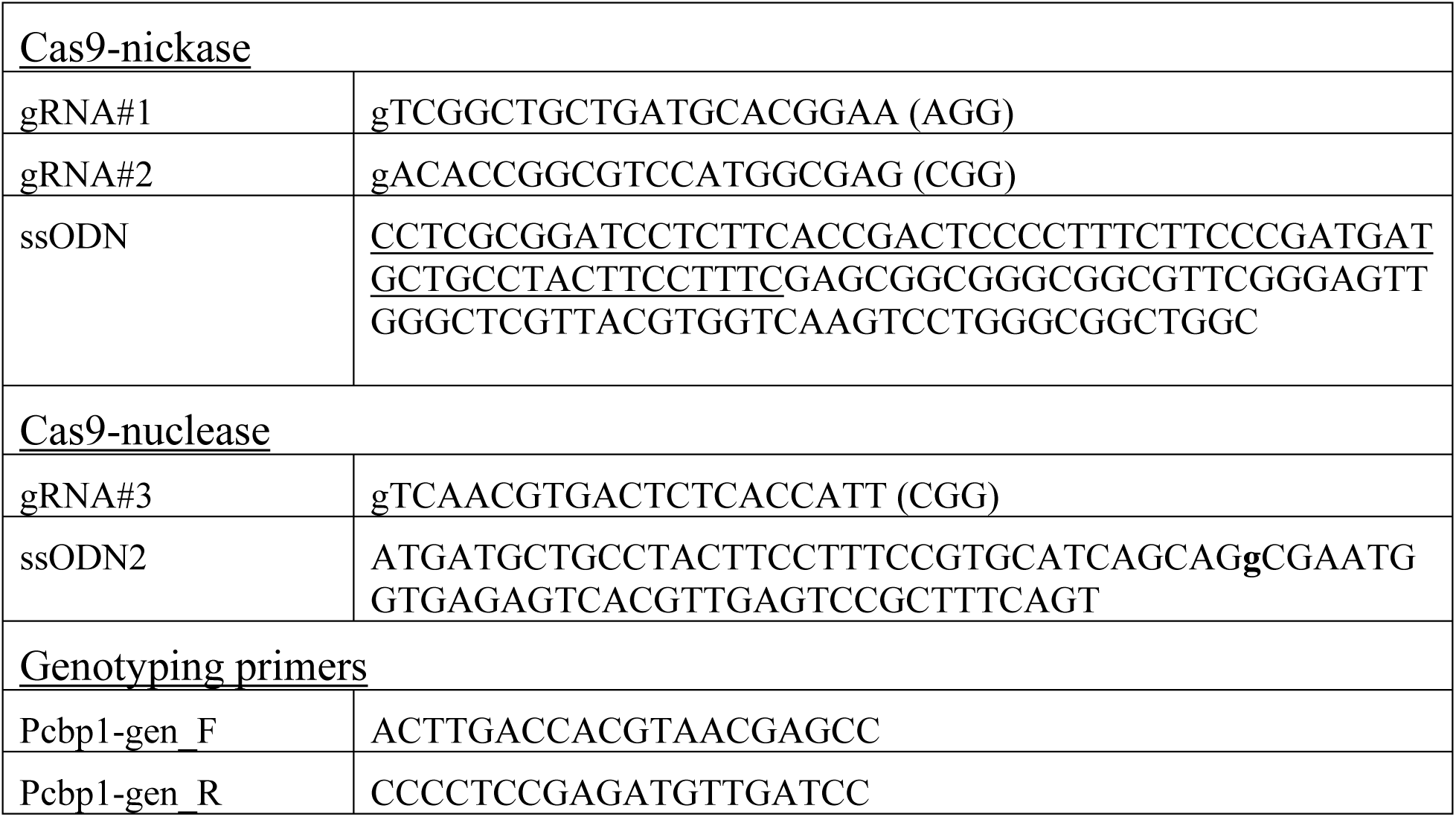
Oligonucleotides used for generation and analysis of *Pcbp1*-heterozygous mice.

### Western blot

For Western blot analysis, cells were harvested and resuspended in PBS. An 1/10 aliquot was taken for protein concentration measurement using the Pierce™ BCA Protein Assay Kit (ThermoFisher Scientific). Then, 2x Laemmli buffer containing β-mercaptoethanol was added, and probes were boiled at 100°C for 5 minutes. For electrophoresis, 10–20 µg of total protein from each probe was loaded onto a 10% acrylamide gel and separated at 80 V. Proteins were transferred onto a nitrocellulose membrane (Bio-Rad), which was then blocked with 5% non-fat milk in PBST. Incubation with primary antibodies was performed in 3% BSA-PBST for 2 h at RT. After 3 or 4 washes in PBST, membranes were incubated with secondary antibodies diluted in 1% fat-free milk in PBST for 1 h, then washed three times in PBST. Chemiluminescence was visualized using the SuperSignal™ West Pico PLUS Chemiluminescent Substrate (ThermoFisher Scientific) and visualized using the ChemiDoc™ Touch Imaging System (Bio-Rad).

### *In vitro* differentiation

Mesodermal and endodermal differentiation was proceeded via embryoid bodies (EB). To this end, 600 Scr or 1200 KO ESCs in 35-µL hanging drops of mES-medium (without LIF) were placed on the Petri dish cover and incubated for five days. Next, EB were transferred to gelatin-coated plastic dishes and cultured for additional 10 days in DMEM/F12 + GlutaMAX supplemented with 10% FBS and 1x Pen-Strep. For neuroectodermal differentiation, Scr and KO EpiSC clumps were plated on fibronectin-coated plastic in N2B27 for three days, then medium was changed to DMEM/F12 + GlutaMAX supplemented with 10% FBS and 1x Pen-Strep for additional three days. After differentiation cells were fixed and subjected to immunocytochemical analysis.

### Cell cycle analysis

Cells were harvested by trypsinization, washed with PBS, and resuspended in 100 µL PBS supplemented with 0.025 % saponin, 50 µg/mL PI, and 0.25 mg/mL RNAse A for 40 minutes at RT. The cells were then analyzed by flow cytometry, and cell cycle distribution was assessed using ModFit LT 3.0 software (Verity Software House, Lexington, MA, USA).

### Time-lapse microscopy

The naïve-to-primed transition was performed as described above except that cells were seeded at a low density (100 cells/cm^2^) for 4 h in the 2i-LIF-N2B27 medium. Then, the medium was replaced with EpiLC medium, and time-lapse microscopy was performed using the CQ1 confocal system (Yokogawa).

### Flow cytometry

Flow cytometry was performed using a CytoFLEX benchtop flow cytometer (Beckman Coulter) with laser wavelengths of 585/42 nm for PI detection (PE) and 660/20 nm for Alexa 647-conjugated secondary antibody detection (APC).

### SeaHorse Energy Profiling

Mitochondrial and glycolytic energy metabolism was assessed using a Seahorse XFe24 Analyzer (Agilent, Santa Clara, CA, USA) and the Seahorse XF Real-Time ATP Rate Assay Kit (Agilent, Santa Clara, CA, USA) according to the manufacturer’s instructions. Cells were seeded in 24-well SeaHorse plates in quadruplicate. Oligomycin and the Rotenone/Antimycin A mixture were added at final concentrations of 3 µM. GlycoATP and mitoATP production were quantified using the SeaHorse XF Real-Time ATP Rate Assay Excel plugin (Agilent, Santa Clara, CA, USA). Data are presented as means ± SEM.

### Quantitative comparison of total protein biosynthesis

The intensity of total protein biosynthesis was quantified using the Protein Synthesis Assay Kit (ab239725) with flow cytometry, following the manufacturer’s instructions. Results are expressed as the means ± SD.

### Surface Sensing of Translation (SUnSET) Assay

The SUnSET assay was performed as described elsewhere [84]. Briefly, ESCs were seeded in 3-cm cell culture dishes either in 2i-LIF-N2B27 or EpiLC medium two days prior to the assay. The cells were then exposed to 10 μg/mL puromycin for 15 minutes. During this period, puromycin, a tyrosyl-tRNA analog, was incorporated into growing polypeptide chains, providing a real-time view of translation levels inside the cells. Following treatment, the cells were harvested and subjected to immunoblotting with anti-puromycin and anti–β-actin antibodies.

### ChIP-seq analysis

Сells were fixed directly on 6-cm culture dishes with 1% formaldehyde, quenched with 0.125 M Glycine, washed in PBS, scraped, and resuspended in sonication buffer containing a protease inhibitor cocktail (PIC, Roche). Chromatin was then fragmented using a CL-18 sonicator (ThermoFisher Scientific) for 10 seconds with 60% amplitude, repeated 10 times with 60-sec cooling on ice after each cycle. Afterwards, immunoprecipitation was performed as follows: chromatin from 1.5x10^6^ ESCs or EpiLCs was incubated with 12 µg of antibodies to Pcbp1, conjugated with MabSelect sure Sepharose (Cytiva). Precipitated IgG-protein-DNA complexes were eluted, and crosslinking was reversed by incubation with proteinase K, NaCl, and EDTA at 55°C for 10 h and at 65°C for 5 h. DNA was purified using phenol:chloroform extraction, dissolved in 50 µL of TE buffer, and used for DNA library preparation and NGS sequencing.

DNA libraries were prepared for sequencing using standard Illumina protocols. The libraries were sequenced to generate high-quality paired-end reads. Raw sequencing reads were processed using fastp [85] to trim adapters and remove low-quality bases with default parameters. The cleaned reads were aligned to the GRCm38/mm10 mouse reference genome using Bowtie2 [86] with default settings. Duplicate reads were identified and marked using Picard MarkDuplicates (http://broadinstitute.github.io/picard/). Peak calling was performed using MACS2 [87] with a stringent false discovery rate threshold (q) < 0.01, using matched input DNA as a control for each corresponding sample.. To generate BigWig files for visualization, MACS2 bdgcmp was used to subtract the bedgraph pileup of the treatment sample from the input control. Peak annotation and enrichment analysis were conducted using the ChIPseeker R package [88] to identify genomic features associated with the called peaks and to gain insights into the biological relevance of the results.

### RNA-seq analysis

Total RNA was extracted using TRIzol (Invitrogen) following the manufacturer’s recommendations. NGS libraries were constructed using the TruSeq Stranded mRNA Kit (Illumina) following the manufacturer’s protocol. 1 μg of total RNA was used as the input material for the protocol. RNA extraction, library construction, and sequencing were performed by Evrogen JSC (Moscow, Russia). RNA-seq libraries were sequenced to generate single-end reads. Raw sequencing reads were processed using fastp [85] to remove adapter sequences and low-quality bases, applying default parameters. Quality control of the cleaned reads was assessed using FastQC [89]. High-quality reads were mapped to the GRCm38/mm10 mouse reference genome using the STAR aligner [90] in two-pass mode to improve splice junction detection.

Further quality control of the aligned reads, including read distribution and gene body coverage, was performed using RSeQC tools [91]. Gene-level read counts were quantified using featureCounts, and differential expression analysis between the knockout (KO) and wild-type (Scr) groups was conducted with DESeq2 [92]. Gene Ontology (GO) enrichment analysis was performed using the FGSEA method, implemented in the clusterProfiler R package [93], to identify enriched biological processes and pathways associated with differentially expressed genes.

### Shotgun proteomics

Cells were washed with PBS and lysed in RIPA buffer (ThermoFisher) supplemented with protease (Roche) and phosphatase inhibitors (Sigma). The samples were subjected to freeze-thaw cycles, sonicated in an ultrasonic bath, and proteins were cleaned from lysis buffer components by acetone precipitation. The protein pellet was resuspended in 8M Urea/50 mM ammonium bicarbonate, and the protein concentration was measured using a Qubit fluorometer (ThermoFisher) with the QuDye Protein Quantification Kit (Lumiprobe) according to the manufacturer’s recommendations. The samples (20 μg) were incubated for 1 h at 37°C in 5 mM DTT (Sigma), followed by incubation in 15 mM iodoacetamide for 30 minutes in the dark at room temperature (Sigma). The samples were then diluted with seven volumes of 50 mM ammonium bicarbonate and incubated for 16 hours at 37°C with 400 ng of trypsin (ratio 1:50; Promega). Finally, the samples were evaporated in Eppendorf Concentrator Plus (Eppendorf), dissolved in water (“LC-MS” grade; LiChrosolv) with 0.1% formic acid (“for LC-MS LiChropur”; Merck), and desalted using C18 Zip Tips (Merck) according to manufacturer’s recommendations. Desalted peptides were evaporated and dissolved in water/0.1% formic acid for further LC-MS/MS analysis.

Approximately 1 μg of each sample was used for shotgun proteomics analysis by HPLC-MS/MS with ion mobility, using a TimsToF Pro mass spectrometer (Bruker Daltonics, Bremen, Germany) coupled with nanoElute UHPLC chromatography (Bruker Daltonics, Bremen, Germany). UHPLC was performed in the two-column separation mode with an Acclaim™ PepMap™ 5-mm Trap Cartridge (Thermo Fisher Scientific, Waltham, MA, USA) and an Aurora Series separation column with nanoZero technology (C18, 25 cm x 75 µm ID, 1.6 µm, C18) in gradient mode at a flow rate of 400 nL/minute and a column temperature of 40°C. Phase A was water/0.1% formic acid, and phase B was acetonitrile/0.1% formic acid (LC-MS Grade). The gradient was from 2% to 35% phase B for 40 minutes, then to 85% phase B for 5 minutes, followed by a wash with 85% phase B for 10 minutes. The column was equilibrated with 4 column volumes before each sample. Electrospray ionization was performed using the CaptiveSpray ion source with a capillary voltage of 1600 V, a 3 L/minute N2 flow, and a source temperature of 180°C. Mass spectrometry acquisition was performed in automatic DDA PASEF mode with a 0.5-sec cycle in positive polarity, fragmenting ions with at least two charges within the m/z range of 100 to 1700 and an ion mobility range of 0.85 to 1.30 1/K0. Each sample was analyzed in at least two analytical replicates.

Protein identification was performed using *Mus musculus* proteins available in UniProt (downloaded 25.02.2022, 34162 proteins), including contaminations. DDA-PASEF data analysis was carried out in FragPipe (v. 17.1) according to the default LFQ-MBR workflow. The search parameters consisted of a parent and fragment mass error tolerance of 10 ppm, a protein and peptide FDR of less than 1%, trypsin as the protease (cleaving after KR, no cleavage before P), and two possible missed cleavage sites. Cysteine carbamidomethylation was set as a fixed modification, while methionine oxidation, STY phosphorylation, and acetylation of protein N-termini were set as variable modifications. Data filtration and imputation of missing values were performed using the NAguideR package for proteomic data analysis [94]. Proteins with missing values in more than half of the samples in at least one biological group and proteins with a coefficient of variation greater than 0.6 were discarded. The optimal method for missing value imputation was selected using “classic criteria,” specifically the “Robust data imputation” approach (“Impseqrob”) [95]. Differential expression analysis was performed by “limma” with Log2-transformed data normalized by quantile normalization [96].

### Metabolomics

Cells cultured in 6-well plates were washed twice with ice-cold PBS, covered with 500 µL of pre-chilled methanol (-80°C), scraped off the plastic, and transferred to Eppendorf tubes. An additional 100 µL of methanol was used to collect the remaining cells. The samples were vortexed and centrifuged for 15 minutes at +4°C. The supernatant (450 µL) was transferred into a new tube for metabolite analysis, while the residual material was used for total protein measurement for subsequent normalization. For derivation of amino acids, 100 µL of the methanol-based total metabolome extract was dried under vacuum (Concentrator Plus; Eppendorf, Germany) at 30°C. The resulting pellet was resuspended in 30 µL of 1-butanol containing 3M hydrochloric acid and incubated at 62°C for 20 minutes with gentle stirring. After cooling for 1-2 minutes, the reaction was stopped by adding 30 µL of deionized water. The samples were dried under vacuum at 30°C, and the resulting pellet was reconstituted in 200 µL of 0.1% formic acid. For derivation of carboxylic acids, 100 µL of the methanol-based total metabolome extract was dried under vacuum at 30°C. The resulting pellet was resuspended in 15 µL of 10 mM triphenylphosphine in acetonitrile to activate carboxylic acids into acyloxyphosphines. Following activation, 15 µL of 10 mM 2-pycolilamine in acetonitrile and 15 µL of 10 mM 2.2’-dithiodipyridine in acetonitrile were added. The reaction mixture was incubated for 20 minutes at 60°C with constant gentle stirring. After the reaction was completed, the samples were cooled down, and the reaction was stopped by adding 45 µL of deionized water. Samples were dried under vacuum at 30°C, and the resulting pellet was reconstituted in 50 µL of 0.1% formic acid.

The analysis was performed using a G6490A triple quadrupole mass spectrometer (Agilent, Inc., Santa Clara, CA, USA). Detection and quantification of amino acids was performed in dynamic SRM (selected reaction monitoring) mode, while carboxylic acids were detected using dynamic neutral loss-SRM (NL-SRM) mode by registering and recording the signal of the 2-pycolylamine–derived product ion (m/z = 109.076). The samples were separated using the UPLC Infinity II 1290 chromatography system (Agilent, Inc., Santa Clara, CA, USA). Samples containing amino acids were loaded at a volume of 5 µL onto the Zorbax Eclipse Plus C18 RRHD column (2.1 × 50 mm, 1.8-µm particle size; Agilent, Inc, Santa Clara, CA, USA) and separated at a flow rate of 0.3 mL/minute in a stepwise gradient of mobile phases A (water) and B (acetonitrile). Samples with carboxylic acids were loaded in a volume of 10 µL and separated at a flow rate of 0.3 mL/minute on an Acquity™ Premier HSS T3 column (2.1 × 150 mm, 1.8 µm particle size; Waters Inc, Ireland) in a gradient of mobile phases A (water) and B (acetonitrile).

### Statistical analysis

Statistical analysis was performed using GraphPad Prism. Unless otherwise specified, data were analyzed using an unpaired Student’s t test. The significance levels were as follows: ns, not significant; *p < 0.05; **p < 0.01; ***p < 0.001; ****p < 0.0001.

## Data availability

The ChIP-seq and RNA-seq datasets have been deposited in the NCBI Gene Expression Omnibus (GEO) database under accession number GSE264037. The mass spectrometry proteomics data have been deposited in the ProteomeXchange Consortium via the PRIDE [97] partner repository with the dataset identifier PXD039287. ChIP-seq, RNA-seq and proteomics data will be available after publication of this study.

The published ChIP-seq datasets used for comparative analysis are: ATAC-seq (GSE155058), H3K27ac (GSE56098), H3K4me3, H3K9me3, and H3K27me3 (GSE155062).

