## Supplementary figures for "Pcbp1 orchestrates amino acid metabolism burst during the naïve-to-primed pluripotency transition"

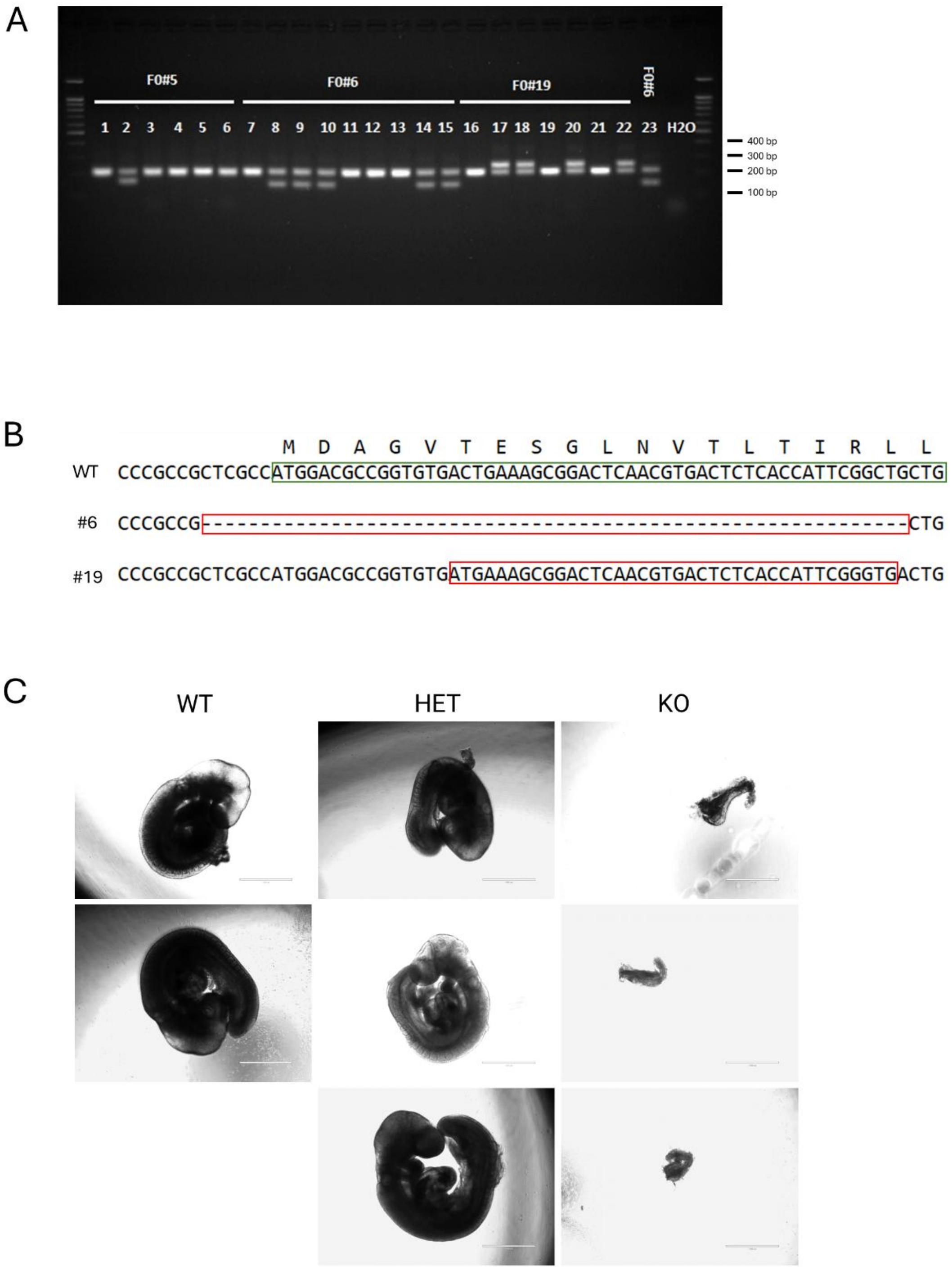


Supplementary Figure 1

(A) Genotyping of the F0 offspring reveals wild-type and heterozygous pups with deletions (#5 and #6) and an insertion #19 at the start of the protein-coding sequence of the Pcbp1 gene. F1 offspring from the #6 and #19 founders were selected for further analysis and mice colony propagation. (B) Alignments showing affected alleles of the #6 (60 bp deletion) and #19 (38 bp insertion) lines. (C) Microphotographs of E9.5 embryos after intercrossing of Pcbp1-heterozygous mice. Scale bar: 1000 μm.

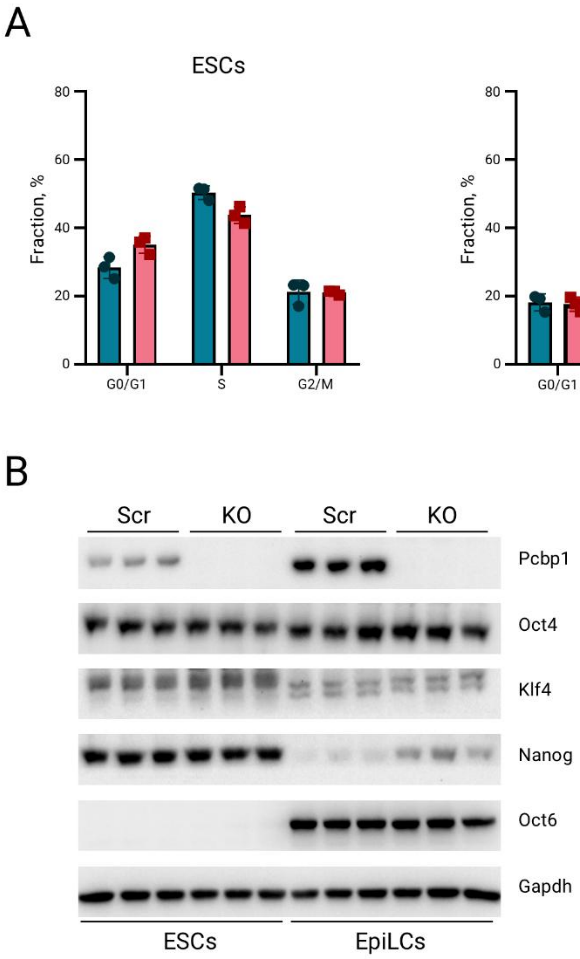


Supplementary Figure 2.

(A) Cell cycle analysis of Scr and KO cells in the naïve (ESC) and formative (EpiLC) states of pluripotency. N = 3 biological replicates (individual cell clones). (B) Western blot analysis of Scr and KO cells in the aforementioned states. N = 3 biological replicates (individual cell clones).

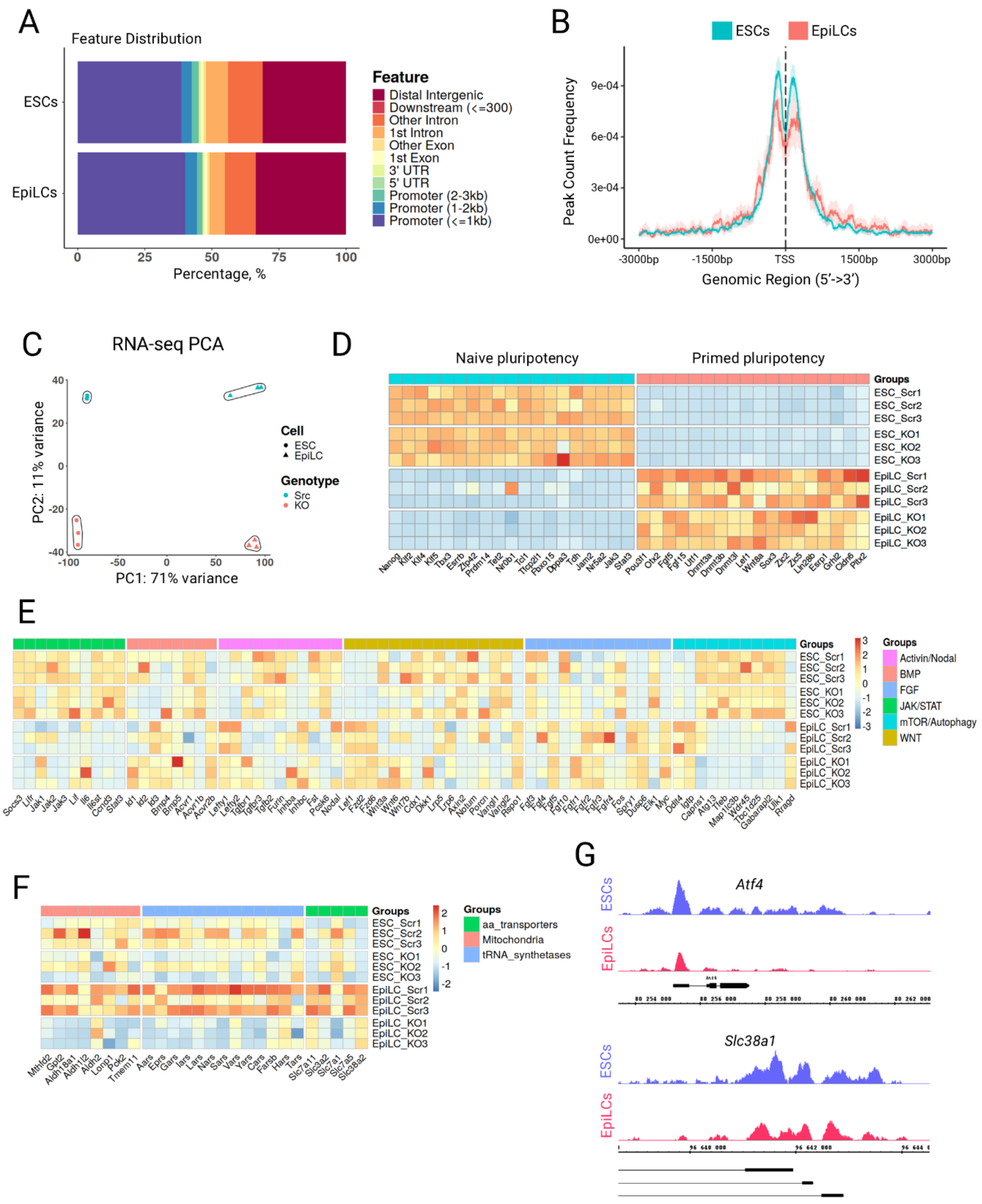

Supplementary Figure 3.

(A) Pcbp1 binding site distribution in ESC and EpiLC genomes revealed by ChIP-seq data analysis. (B) Pcbp1 binding site distribution around transcription start sites (TSS). (C) Principal component analysis showing clustering of RNA-seq samples by Scr vs. KO and ESC vs. EpiLC. (D) Heatmap displaying gene expression profiles related to naïve and primed pluripotency in Scr and KO cells (N = 3 biological replicates, i.e. individual cell clones). (E) Heatmap showing expression of genes related to different signaling pathways typical for pluripotent stem cells. (F) Heatmap showing expression of typical targets of the transcription factor Atf4. (G) Integrated genome browser snapshots of ChIP-seq data showing examples of genes occupied by Pcbp1 in ESCs and EpiLCs.

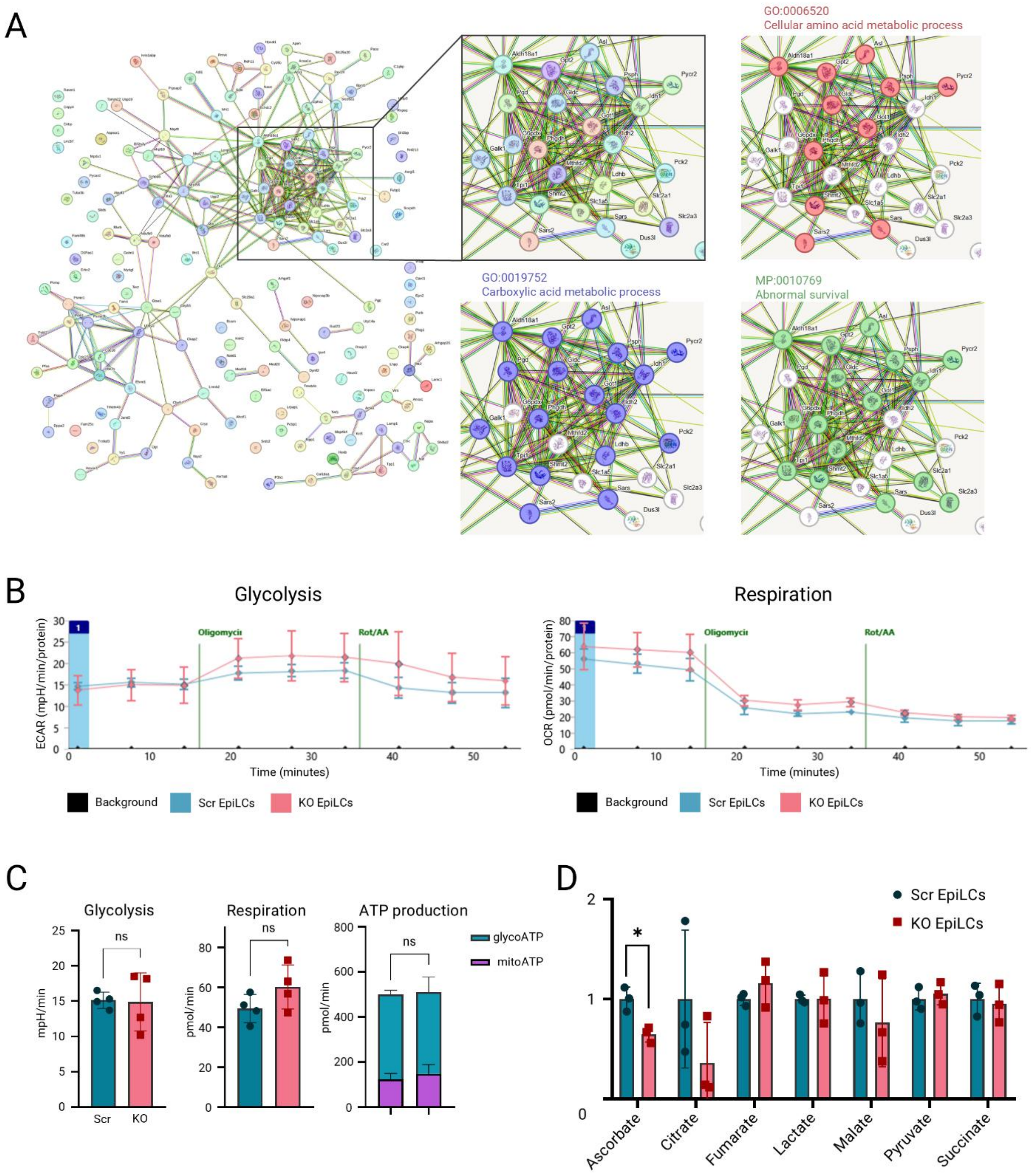


Supplementary Figure 4.

(A) Clustering of differentially presented proteins in EpiLCs using STRING (string-db.org), with several enriched pathways highlighted. (B) ECAR (glycolysis) and OCR (respiration) measurements using the SeaHorse XFe96 analyzer. (C) Energy metabolism of Scr and KO EpiLCs using a SeaHorse analyzer, measuring glycolysis (ECAR), respiration (OCR), and calculated ATP production (N = 4 technical replicates). (D) Metabolome analysis of carboxylic acids in Scr and KO EpiLCs. N = 3 biological replicates (individual clones); *p<0.05.
